# Policy optimization emerges from noisy representation learning

**DOI:** 10.1101/2024.11.01.621621

**Authors:** Jonah W. Brenner, Chenguang Li, Gabriel Kreiman

**Affiliations:** Department of Physics, Harvard University, Cambridge, Massachusetts 02138, USA; Department of Applied Physics, Stanford University, Stanford, California 94305, USA; Sainsbury Wellcome Centre, University College London, London W1T 4JG, UK; Boston Children’s Hospital, Harvard Medical School, Boston, Massachusetts 02115, USA

## Abstract

Biological nervous systems learn both internal representations of the world and behavioral policies for acting within it. Motivated by growing evidence that representation learning is a fundamental principle underlying synaptic plasticity, we introduce **N**eural **S**tochastic **M**odulation (NSM): a theory of learning in which policy optimization emerges from reward-modulated noise layered on top of plasticity rules designed for representation learning. In NSM, reward-modulated noise shapes the steady-state weight distribution, guiding the network toward solutions that capture meaningful features while also maximizing reward. Interestingly, the evolving internal representations produced by our model mirror neural coding changes observed experimentally during task learning. Our results suggest that reward-modulated noise can serve as a minimal and biologically plausible mechanism for integrating representation and policy learning in the brain.

## I. INTRODUCTION

Biological nervous systems learn internal representations—structured neural encodings that capture or predict relevant features of the world—and behavioral policies that guide action within their environments [1–3]. These capacities are thought to emerge primarily from synaptic plasticity: the process by pairs of neurons adapt their connection strengths so that the macroscopic network performs different cognitive functions [4, 5]. A central goal of theoretical neuroscience is to identify classes of plasticity rules that both account for the biologically observed features of synaptic plasticity, and construct networks that extract structure from sensory inputs and act to maximize reward.

A growing body of experimental and computational work suggests that plasticity rules derived from unsupervised or self-supervised representation learning objectives account for many observed features of synaptic plasticity, for instance in the visual system [6–11], in hippocampus [12–14], and in neocortex [15]. Motivated by these successes, we take this idea to its extreme and ask: What if, as a zeroth-order appenximation, all of the brain’s deterministic plasticity rules were solely dedicated to representation learning? Could such a system not only build internal models of the world but also learn how to act within it?

This article introduces **N**eural **S**tochastic **M**odulation (NSM), a framework for how policy optimization can emerge from noisy representation learning in the nervous system. NSM treats the brain’s intrinsic noise as a functional component of learning. It hypothesizes that neuromodulation makes the nervous system’s noise reward-dependent, decreasing its magnitude when the weight configuration yields a rewarding policy. This “stochastic modulation” shapes the stationary distribution over the network weights, steering the network toward rewarding policies even though its deterministic plasticity rules optimize a reward-agnostic objective.

We present general analytical results showing that the NSM framework enables agents to learn rewarding policies. We illustrate these results using an agent with a one-layer network operating under a simple, linear representation learning objective. We evaluate the agent’s behavior in two tasks. In the first, the features extracted by the representation learning objective fully capture the task-relevant information—meaning that optimal policies lie within the solution space of the objective. Using analytical arguments and simulations, we demonstrate that the agent is guaranteed to discover an optimal policy by exploring the space of representations consistent with its objective. In the second task, solving the problem requires learning features that deviate from the original representation learning objective. In this setting, we show that reward-dependent noise drives the agent to minimize an effective loss that explicitly balances normative representation learning with reward, enabling successful policy learning. Taken together, these results suggest that modest, biologically plausible extensions of representation learning rules can support emergent policy optimization, offering a unifying perspective on how representation learning and behavior might co-emerge in the brain.

## II. RELATED WORK

Theoretical neuroscience has long sought to understand the synaptic plasticity rules by which biological neural networks learn to enact effective policies. Early models addressed this by proposing reward-modulated spike-timing dependent plasticity (STDP) as a mechanism for reinforcement learning, demonstrating how local plasticity paired with a global reward signal can support synaptic updates that improve behavior [16, 17]. These models remain influential due to their biological plausibility: neuromodulatory systems are conserved across species—from *C. elegans* to humans—and observed STDP dynamics align across many organisms [18–21]. However, networks relying solely on this paradigm typically solve only very simple tasks, such as discriminating spike trains.

Crucially, these early models focused on directly learning reward-driven behavior, without considering representation learning as an independent computational goal. In contrast, representation learning has proven essential in artificial agents, where objectives like compression and prediction greatly enhance performance in complex tasks [22, 23]. Accumulating evidence also suggests that biological neural systems continually perform representation learning as an independent computational goal, even when rewards are sparse or absent [24, 25]. These findings call for a unified view in which policy optimization arises as an emergent property of networks focused on building useful internal representations.

Our work takes a step in this direction. Like a recent biologically-plausible plasticity model that showed success solving Atari games with spiking networks [26], we base plasticity on representation learning rules. However, unlike that approach—which builds an additional policy-learning system on top—we propose that modulation of intrinsic synaptic noise can steer the network toward rewarding configurations. The use of noise as a computational tool is not new [27–29]. But, to our knowledge, the key idea that reward-dependent noise can bias the steady-state distribution of a network, trained on a reward-agnostic objective, towards rewarding solutions has not yet been explored.

## III. THE NEURAL STOCHASTIC MODULATION FRAMEWORK

We now introduce **N**eural **S**tochastic **M**odulation (NSM): a framework in which a single noisy neural network both learns representations and controls actions, while a separate modulatory system tunes its synaptic noise to facilitate policy optimization.

The NSM framework has three key components:

1. A neural network that is designed for representation learning,
2. A motor function that maps neural activity to actions, and
3. A modulatory system that shapes synaptic noise to facilitate policy optimization.

Each component is described below and illustrated in Figure 1. We assume throughout that our agent operates within a Markov Decision Process (MDP), which is a natural formalism for modeling a biological organism interacting in an environment.^1^

**FIG. 1.**
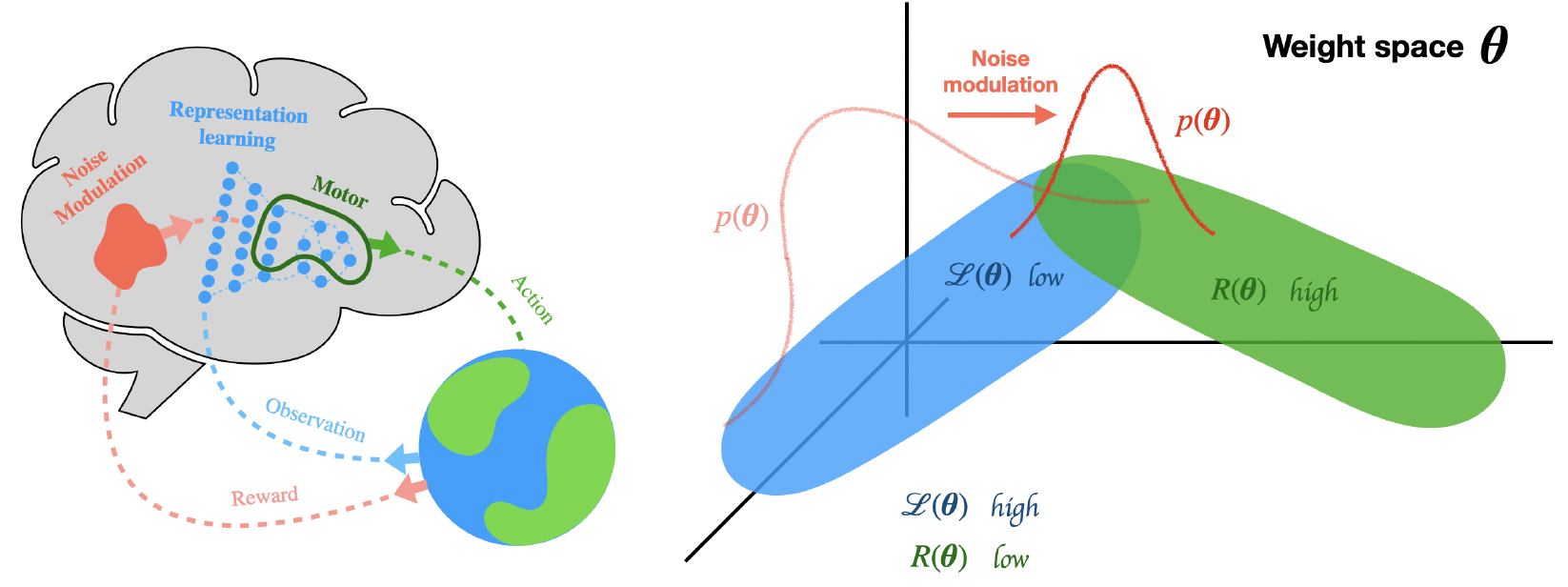
**(Left)** Schematic of the Neural Stochastic Modulation (NSM) framework. A neural network designed for representation learning (blue) receives observations from the environment and extracts features. A motor function (green) maps the network’s activity to an action, defining the agent’s policy. A modulatory system (red) adjusts the level of intrinsic noise in the weight dynamics based on an estimate of the policy’s reward. **(Right)** Schematic of how reward-dependent noise modulation facilitates policy optimization. The stationary distribution over weights, *p*(***θ***), concentrates in regions where the representation learning objective ℒ (***θ***) is low (blue) when there is no noise modulation (light red). Introducing modulation shifts this distribution (dark red) toward regions where both ℒ (***θ***) is low and the policy reward *R*(***θ***) is high (green).

### A. A representation learning objective shapes the network’s weight dynamics

Consistent with biology, we consider an agent that behaves and learns online, continuously adjusting its neural network’s activities and weights as it interacts with the environment. We begin by describing the dynamics of the agent’s neural network activities and weights.

Let the network be parameterized by synaptic weights ***θ*** and neural activities **x** ∈ ℝ^*N*^, where *N* is the number of neurons. The network can have any architecture. The choice of activity dynamics is not the focus of NSM, but for concreteness, we assume that they are stochastic and first-order in time:

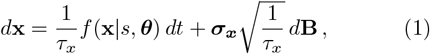

where *d***B** is a standard Wiener process, *τ*_*x*_ is the timescale of the activity dynamics, and ***σ***_***x***_ shapes the statistics of the activity noise. The function *f* (**x** | *s*, ***θ***) ∈ ℝ^*N*^ defines the baseline neural activity dynamics without noise in the environmental state *s*. We express dynamics in terms of timescales *τ* rather than learning rates *η* = 1*/τ* throughout this work to emphasize that we are modeling biological systems, not engineering learning algorithms. Our results do not rely on this specific form of activity dynamics.

We now turn to weight dynamics, the focus of NSM. The framework postulates that weights perform noisy gradient descent on a representation learning objective ℒ (**x**, *s*, ***θ***). This objective encourages the network to learn meaningful representations of the environmental inputs. For example, it may encourage the network to extract features of the MDP states or their transition dynamics. We refer to the resulting representations as *normative* because they are dictated by the optimization of ℒ, independent of task reward. The weight dynamics are:

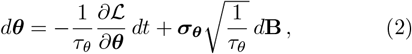

where the notation follows Eq. (1) and the subscripts signify that the parameters of the activity and weight dynamics can differ. Conceptually, we include timescales for both activity and weight dynamics (rather than just one) to allow for comparison with a third relevant timescale in the problem: the timescale of external environmental dynamics.

Often one assumes *τ*_*x*_ ≪ *τ*_*θ*_, such that activity dynamics evolve on a faster timescale than weight updates. As a result, the network first responds to an environmental state *s* by adjusting its activities **x**, and subsequently modifies its synaptic weights ***θ*** based on those activity patterns.

#### Example Agent

To illustrate the NSM framework, we introduce a simple example agent. This minimal agent captures key features of NSM while remaining analytically tractable and easily interpretable. We will return to this example as a reference throughout the paper.

The agent’s representation learning network optimizes an online formulation of the Similarity Matching (SM) objective, originally introduced in [30]. SM encourages the network to project its inputs onto their principal subspace, a form of dimensionality reduction. The objective also gives rise to Hebbian plasticity, which emphasizes the biological plausibility of our framework.

We let the network have one layer, with neural activities **x** ∈ ℝ^2^ and weights ***θ*** = (**W**_0_, **W**_1_). The matrix **W**_0_ ∈ ℝ^2*×D*^ represents feedforward connections from *D*dimensional input states **s** ∈ ℝ^*D*^, while **W**_1_ ∈ ℝ^2*×*2^ represents recurrent lateral (inhibitory) connections within the network.^2^

The online SM objective for this architecture is:

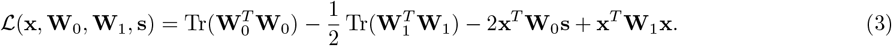

Following the NSM prescription for weight dynamics, and in this example, letting the activity dynamics also perform gradient descent on ℒ, we give the network dynamics the form:

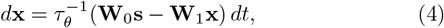

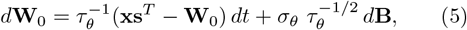

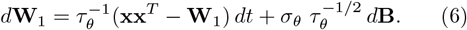

For simplicity, we set the activity noise to zero to make the network’s activity dynamics deterministic. We will see soon that this also makes the agent’s policy deterministic, which will simplify future analysis. We also let ***σ***_***θ***_ be diagonal with equal elements *σ*_*θ*_. At each timestep of the MDP, the activities evolve under the differential equation (4) until they converge, followed by a discrete weight update.

### B. A motor function defines the agent’s policy

Under NSM, the same neural network simultaneously learns representations and governs action selection. We now introduce the motor function, which maps the network’s activities to an action and thereby defines the agent’s policy.

Let *M* : ℝ^*N*^ → 𝒜 be a motor function that maps neural activities **x** to an action *a* ∈ 𝒜. We assume *M* is time-invariant.^3^ Biologically, we can imagine that this motor function arises due to projections from the neural activities **x** to a motor control system or from movements of the muscles directly controlled by **x**.

This motor function defines the agent’s policy, the distribution of possible actions *a* that it could take in response to a state *s*. In the general case of noisy activity dynamics, a distribution *g*_*θ*_(**x** | *s*) describes the possible network responses to a state *s*. The policy induced by the motor function is given by the distribution of the actions associated with these network responses

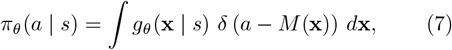

where *δ*(·) denotes the Dirac delta function. This relation holds for discrete and continuous actions.

#### Example Agent

The example agent has the action space 𝒜 = {−1, 1}. We define a simple linear decoding rule:

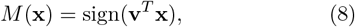

where **v**^*T*^ = ( −1, 1). This defines a linear decision boundary in representation (activity) space. Notice that our agent’s policy is deterministic. Because we set its activity noise to zero, it maps a state *s* deterministically to an activity pattern, which in turn defines an action through *M*.

### C. Modulating the stochastic weight dynamics facilitates policy optimization

The central postulate of NSM is that an additional system modulates the noise statistics of the network. In this work, we focus on the modulation of weight noise statistics, though Appendix H argues that similar results can hold for activity modulation.

The weight noise in Eq. (2) depends on two parameters: the noise matrix ***σ***_***θ***_ and the timescale *τ*_*θ*_. NSM proposes that reward-dependent modulation acts through one or both. We refer to changes in ***σ***_***θ***_ as *direct noise modulation*, since they alter the noise explicitly, and changes in *τ*_*θ*_ as *indirect noise modulation*, since they affect the noise via the timescale. In both cases, policy optimization emerges from reward-dependent noise, even though the expected value of the weight updates follow only the gradient of the representation learning objective.

#### Direct noise modulation

Here, the modulatory system adjusts the weight noise matrix according to an estimate of the expected reward *R*:

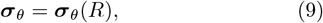

where the eigenvalues of ***σ***_*θ*_(*R*) decrease monotonically with *R*. This mechanism reshapes the stationary distribution over the weights by “cooling” the system when the agent discovers rewarding policies. The reward-dependent noise often acts like a regularizer, encouraging the network to favor solutions to ℒ that also achieve high task reward.

To make this idea concrete, let us assume that the noise magnitude is small enough that the weights remain approximately confined to the solution space of ℒ. Let ***σ***_***θ***_(R) be diagonal with elements *σ*_*θ*_(*R*), where

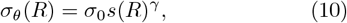

and *s*(*R*) is a monotonically decreasing function of *R, σ*_0_ *>* 0 is the baseline noise level, and *γ >* 0 is a scalar whose meaning will soon become clear. With this choice, the stationary distribution of weights restricted to the solution space of ℒ becomes

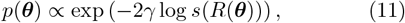

as derived in Appendix C.

This distribution is equivalent to that of a system performing gradient descent on the effective objective

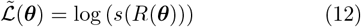

under fixed synaptic noise *γ*^−1*/*2^ within this solution space. Since *s*(*R*) decreases with reward, minimizing 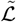 is equivalent to maximizing *R*. Like a regularizer, direct noise modulation steers the system towards high-reward weight configurations within the solution space of the representation learning objective.

While direct modulation noise modulation makes reward a regularizer on representation learning, indirect modulation instead creates an explicit trade-off between representation learning and reward optimization.

#### Indirect noise modulation

The modulatory system can also control the network’s synaptic noise through the timescale of the stochastic weight dynamics:

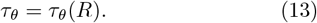

This mechanism can explicitly balance representation learning with reward optimization. In particular, if we choose the timescale modulation to take the form:

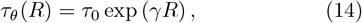

with *τ*_0_ *>* 0, and let the noise matrix ***σ***_***θ***_ be diagonal with elements *σ*_*θ*_, then the stationary distribution over the weights becomes:

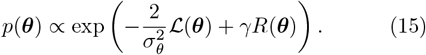

We derive this result in Appendix D. Remarkably, this is equivalent to the stationary distribution that arises from explicit gradient descent on the effective objective:

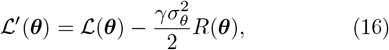

under fixed synaptic noise *σ*_*θ*_. With respect to the stationary distribution of learning, the weight dynamics in Eq. (2) with indirect noise modulation are equivalent to explicit gradient optimization of the reward function and representation learning objective together. To our knowledge, this is a new result in the context of learning.^4^ Note here that even though the timescale is being modulated, the noise is still the crucial ingredient: If *σ*_*θ*_ → 0, the reward drops out of the effective loss.

We emphasize that neither the direct nor the indirect modulation results assume a specific network architecture or a representation learning objective ℒ. Therefore, these results hold across agents across different architectures and representation learning objectives. While we focus on the example agent in the main text, we present numerical support for this generality in Appendix F.

#### Example Agent

We will demonstrate direct noise modulation Eq. (9) in the first task that we consider and indirect noise modulation Eq. (14) in the second.

In the first task, we let ***σ***_*θ*_ be diagonal with elements:^5^

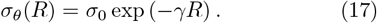

This exponential form corresponds to choosing *s*(−*R*) = exp(*R*) in Eq. (10). We adopt this form for clarity, as it allows a straightforward comparison with the formulation of indirect modulation [Eq. (14)].

In the second task, the noise amplitude *σ*_*θ*_ is held fixed, while the weight timescale *τ*_*θ*_(*R*) is modulated according to Eq. (14).

While this work focuses on NSM in the context of policy optimization emerging from networks designed for representation learning, the framework is more general and can be applied more broadly. We outline this generality in Appendix B.

## IV. RESULTS

### A. Direct noise modulation biases learning towards rewarding policies within the solution space of ℒ

Direct noise modulation encourages the agent to explore weight configurations in and around the solution space of its representation objective ℒ in search of rewarding policies. When ℒ captures all task-relevant structure, policy learning reduces to selecting among equivalent representations within this space.

We analyze this setting analytically by studying the behavior of our example agent ℒ in a contextual bandit task. We compute the steady-state distribution of the network weights in the presence of direct noise modulation. We show that the distribution is biased so that the agent is guaranteed to find a rewarding policy as time goes to infinity.^6^ This illustrates NSM’s core principle: reward, acting only through noise modulation, can guide an agent to learn effective policies.

#### 1. The contextual bandit task

The contextual bandit is a special case of the MDP in which state transitions are independent of actions. In our setup, each 10-dimensional input **s** is generated by multiplying a Gaussian vector by a random sign:

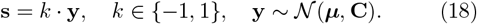

The covariance matrix **C** has eigenvalues *λ*_1_ = 0.1 and *λ*_2:10_ = 0.05. The mean ***µ*** is aligned with the second principal component of **C** and scaled to norm ∥***µ***∥ = 2. This generates inputs from two linearly separable Gaussian clusters.

At each timestep, the agent observes **s** and selects an action *a* ∈ {−1, 1}. It receives a reward if its action corresponds to the random sign, i.e., *a* = *k*. Figure 2 illustrates the setup.

**FIG. 2.**
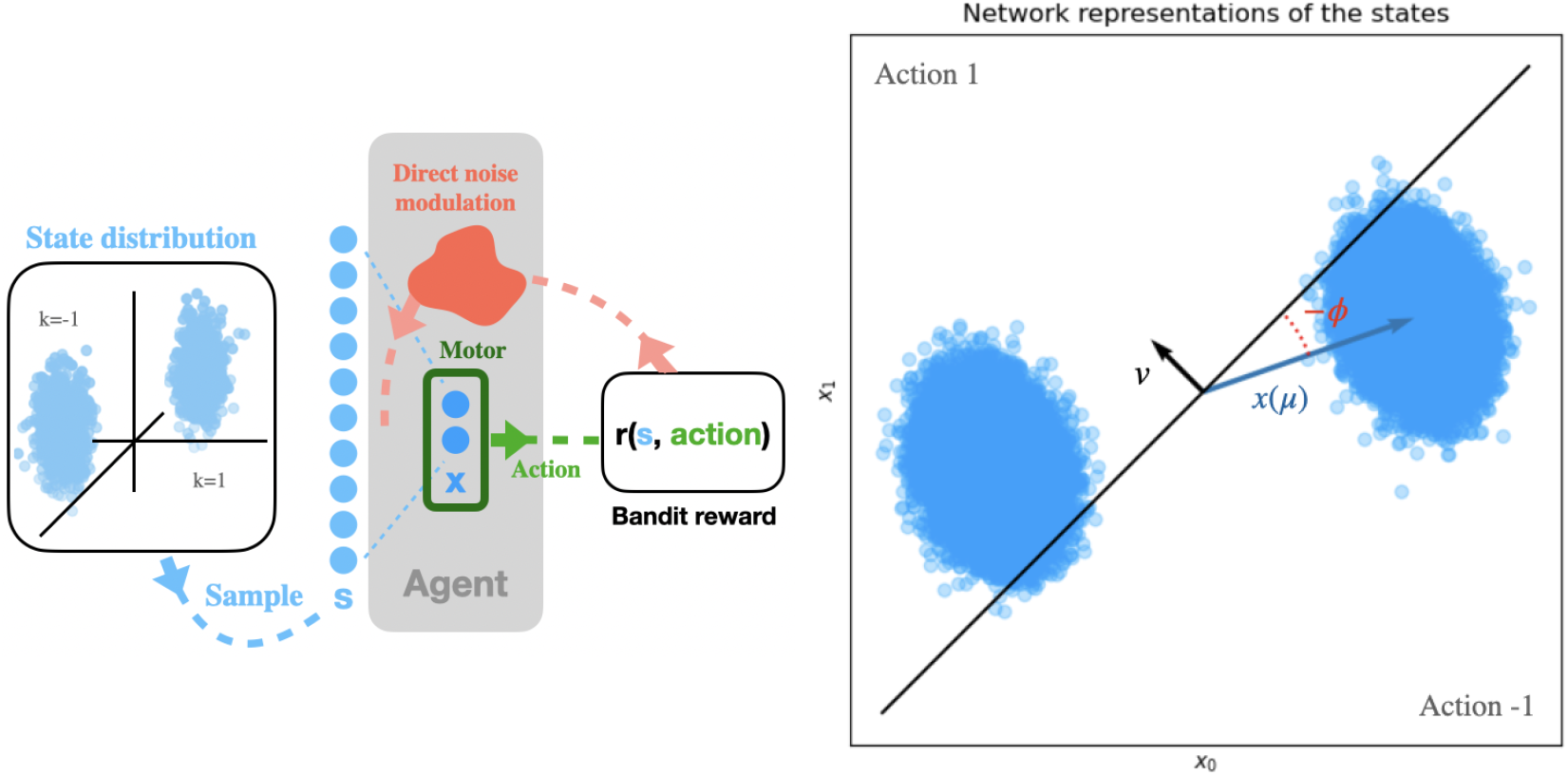
**(Left)** Schematic of the agent in the contextual bandit task. The agent receives a state **s** (light blue), and its activities respond, resulting in an action. The neuromodulatory system receives the reward associated with this action and adjusts the weight learning rate. The weights then update according to ℒ, and the task continues. The agent’s goal is to take the action corresponding to the random sign *k*. **(Right)** Illustration of all relevant variables in the network’s representation space, where the axes denote the activity of its two neurons. Royal blue points show the network representations of the states **s**, denoted **x**(**s**). The vector **v** defines a classifier that maps these representations to the action 1 or −1. Finally, *ϕ* tracks the configuration of the network weights through policy learning.

#### 2. A order single parameter ϕ describes the network’s weight configuration

Our goal is to show that the agent learns a rewarding policy given strong enough direct noise modulation. We begin by showing that the network’s weight configuration during learning can be characterized by a single order parameter, *ϕ*. This simplification is important because it allows us to analyze which policies the agent learns by studying the steady-state distribution over *ϕ*.

To understand where *ϕ* comes from, we examine the solution space of the network’s representation objective. This objective encourages the network to project inputs **s** onto their top two principal components. As noted in Ref. [32], the loss ℒ remains invariant under the family of weight transformations of the form:

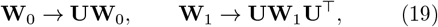

which results in the network representations transforming as:

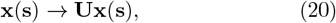

where **U**^⊤^**U** = 𝕀. Because the network representations **x**(**s**) lie in ℝ^2^, the allowed transformations **U** are two-dimensional rotations. So, each matrix **U** can be parameterized by a single angle *ϕ*. Therefore, there is a one-to-one correspondence between the set of weight configurations in the solution space of ℒ and the values of *ϕ* ∈ [0, 2*π*).

Conveniently, the angle *ϕ* also determines the rotational orientation of the network’s representations. Therefore, *ϕ* determines the agent’s policy (see Fig. 2). We choose *ϕ* = 0 to be the reference orientation where the network’s representation of the input mean, **x**(***µ***), is aligned with the decision boundary used by the agent to determine its actions. Notice that different values of *ϕ* correspond to different policies. For example, 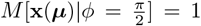 whereas 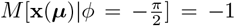. This means that different values of *ϕ* will also correspond to different average rewards.

We can describe the evolution of the network weights during learning with *ϕ* as long as the weights remain in the solution space of ℒ. This assumption does not hold in general: as we discuss later, reward-dependent noise can stabilize points outside the solution space. However, in this task, we find empirically (see Appendix G) that the representation learning loss decreases rapidly and remains near its minimum throughout training. This occurs because the inputs’ principal subspace captures all task-relevant information. We can safely assume that, after a brief convergence period, the network remains approximately within the solution space of ℒ.

Thus, from a steady-state perspective, the state of the network can be fully described by *ϕ*. In what follows, we model the network’s long-time learning as stochastic dynamics for *ϕ*, and compute its limiting distribution in the presence of reward-modulated noise.

#### 3. Reward-dependent noise biases the network towards rewarding policies

Random weight transformations of the form (19), where **U** represents an infinitesimal rotation, cause the network weights to explore the solution space of ℒ. As a result, the network representations rotate, and *ϕ* follows stochastic dynamics with reward-dependent noise. The noise is small when *ϕ* corresponds to a network that follows a rewarding policy and large otherwise.

The stationary distribution of *ϕ* reveals which policies the network tends to learn. Because the weights remain approximately confined to the solution space of the representation learning objective, we can use the earlier result (Eq. 11), which gives the stationary distribution *p*(***θ***) under direct noise modulation within this region. Let ℛ denote this solution space. Within ℛ, the network weights ***θ*** are uniquely determined by the angle *ϕ*, such that ***θ***(*ϕ*) = *h*(*ϕ*) for some function *h* (see Eq. 19). This function is one-to-one, and we denote its inverse by *h*^−1^(***θ***). The stationary distribution over 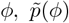, can therefore be written as

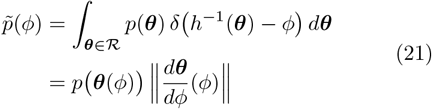

where 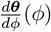 is a vector containing the derivative of each weight with respect to *ϕ*, and ∥.∥ denotes the L2 norm. We show in Appendix E that the Jacobian factor 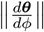 is constant and can therefore be absorbed into the normalization of the distribution.

Substituting the specific noise modulation rule used in this example (Eq. 17) into the stationary weight distribution *p*(***θ***) from Eq. (11) gives

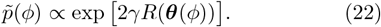

To make this expression explicit, we derive in Appendix K an approximate analytical form for the expected reward *R* as a function of *ϕ* in the contextual bandit task:

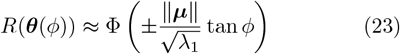

where F is the standard normal CDF, and *λ*_1_ is the largest eigenvalue of **C**. We take the negative sign when 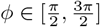 and the positive sign otherwise.

Substituting Eq. (23) into the stationary distribution yields an explicit form for the limiting distribution of *ϕ*:

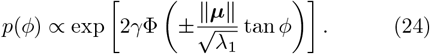

We can understand this expression as a distribution over policies, as *ϕ* determines the agent’s policy. We can also use this expression to find the expected reward received by an agent ·[*R*], which we compute numerically using Eq. (23) and Eq. (24). In Figure 3, we see that as *γ* increases, ·[*R*] → 1. With strong modulation, the agent is guaranteed to learn a policy that maximizes reward.

**FIG. 3.**
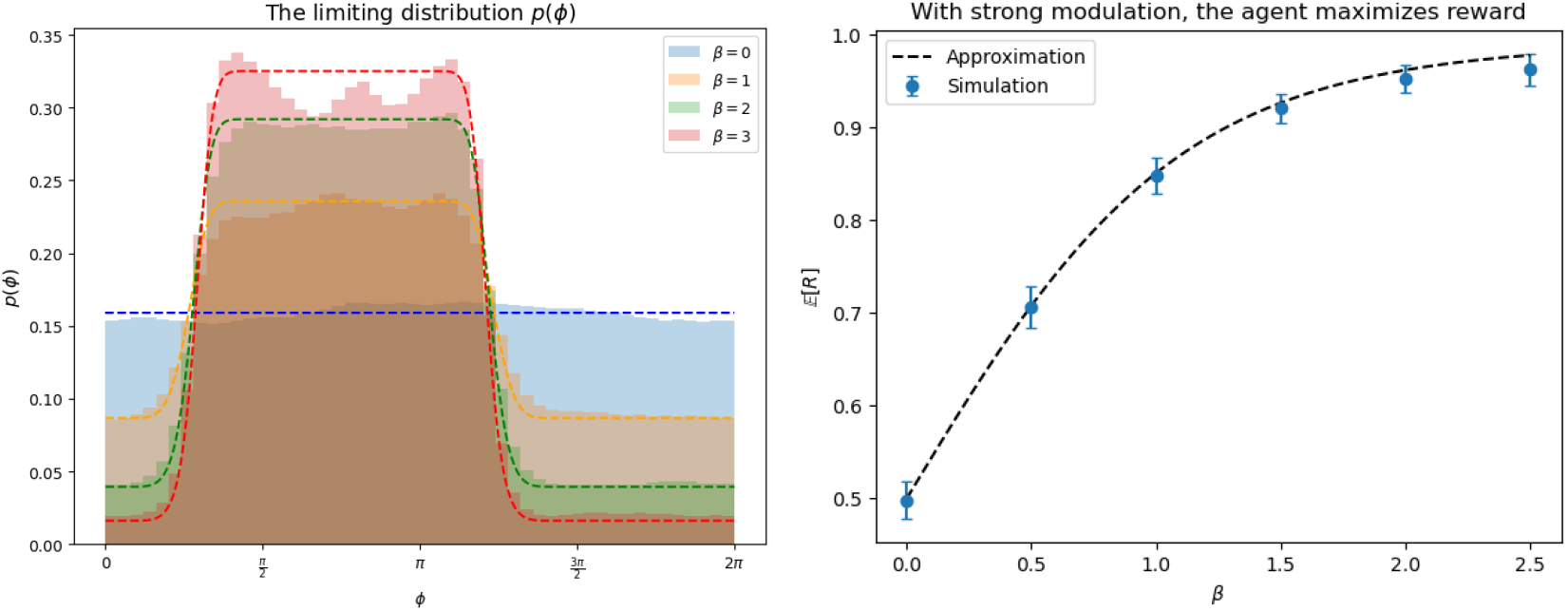
**(Left)** The limiting distribution of *ϕ*. The dashed lines denote our approximations for different *γ*, and the histograms show the simulation data. With *γ* = 0 (no noise modulation), *ϕ* is uniformly distributed in the solution space of ℒ. With larger *γ*, the agent becomes biased towards rewarding angles. Data generated from 100 seeds of 500,000 timesteps. Error bars denote standard deviation across seeds. **(Right)** The expected reward received by the agent in the limit that time goes to infinity. The agent is guaranteed to maximize reward for large *γ* (stronger modulation). Our approximation agrees with simulations.

Policy optimization in this setting emerges entirely through reward-modulated noise, despite the network optimizing a reward-independent objective. From a steady-state perspective, the noise biases the weight distribution toward rewarding configurations within the solution space of ℒ. From a dynamical perspective, it facilitates a diffusive search over equivalent representations of the stimuli, increasing the probability of settling in those that yield high reward.

### B. Indirect noise modulation balances normative representation learning with reward by minimizing ℒ′

We now consider what happens when the representation learning objective does ℒ *not* capture all task-relevant information—that is, when no optimal policy exists within its solution space. Recent work in the context of stochastic gradient descent found that the weight noise matrix ***σ***_*θ*_(***θ***) and the loss ℒ (***θ***) jointly determine the attractive points in weight configuration space [33].

So, directly modulating the noise can bias the system toward policies that improve reward, even at the cost of deviating from normative representations.

However, indirect noise modulation offers a more principled mechanism. As shown in Eq. (16), it causes the network to minimize an *effective* loss ℒ′, which combines representation learning and reward into a single objective. When reward and ℒ are misaligned, minimizing ℒ′ allows the network to trade off between them explicitly.

To demonstrate this, we turn to Cart Pole, a classic control task. The agent’s goal is to balance a pole on a moving cart by pushing it left or right at each timestep. The task is simple enough to be solved by our example agent. But crucially, projecting state observations onto their two-dimensional principal subspace does not preserve enough information to support a good policy (see Appendix L). As a result, learning a successful policy requires deviating from the normative representations that minimize ℒ.

Figure 4 (left) shows that the agent diverges from the solution space of ℒ in order to reduce its effective loss ℒ′. By accepting a higher representation loss, the network extracts additional task-relevant features that improve action selection. Through this process, the system balances explicitly normative representation learning with maximizing reward and solves the task.

**FIG. 4.**
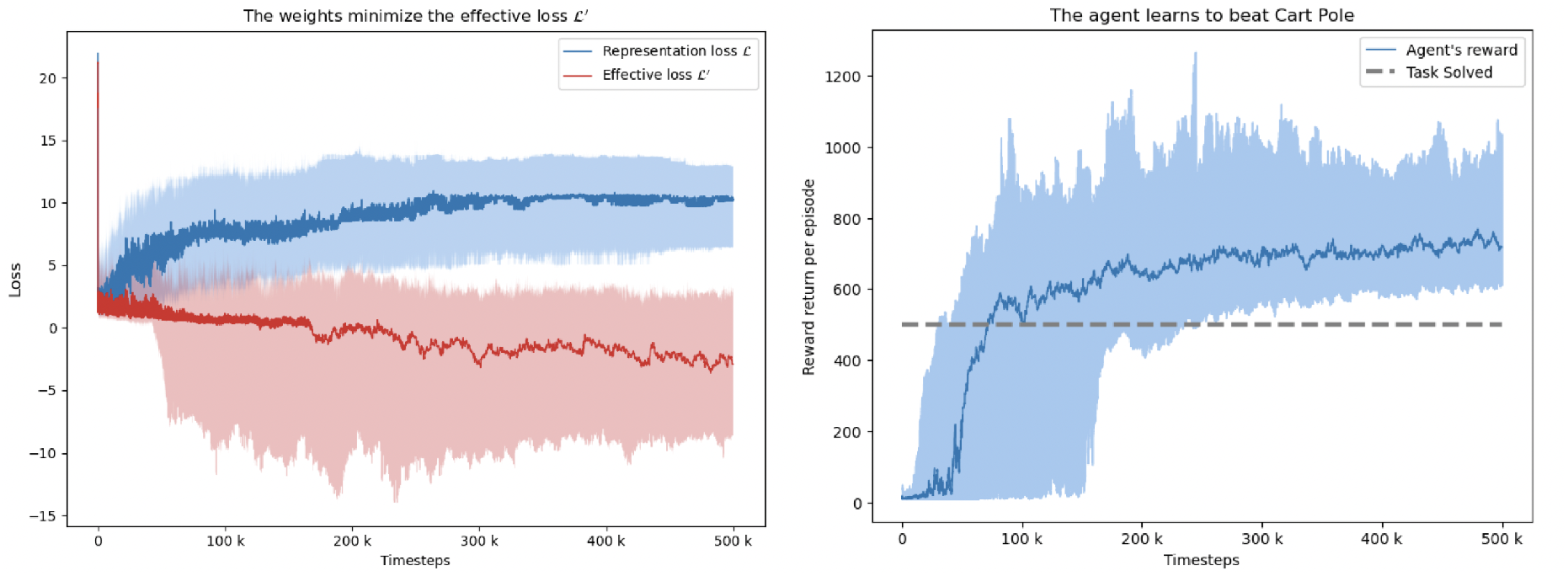
**(Left)** The representation loss ℒ and the effective loss ℒ′ over time in the Cart Pole task. The network’s steady-state distribution minimizes ℒ′, not ℒ. **(Right)** The agent learns to beat Cart Pole. In both plots, curves show the median and middle 50% over 100 seeds plotted through 500,000 timesteps of interaction.

Through the modulation of noise, NSM provides a minimal and biologically plausible mechanism by which representation and policy learning can emerge from a unified process. This raises a natural next question: to what extent is NSM consistent with experimental observations?

## V. CONSISTENCY WITH EXPERIMENTAL OBSERVATIONS

While NSM is a general framework, we focus on the case where ℒ reflects a representation learning objective and *R* denotes reward. In this setting, NSM qualitatively reproduces key phenomena observed during task learning. We highlight two examples: representation drift and neural reassociation.

Representation drift describes the slow change in the neural codes of stimuli over time. Crucially, these changes maintain the information encoded in the neural population [34–36]. Representation drift emerges naturally from NSM when ℒ is a representation learning objective. We saw representation drift clearly in the contextual bandit task: the network responses rotated while maintaining their population structure. This result has been noted before in [32]. NSM suggests a potential feature of this drift: It can aid task learning by sampling behavioral policies, even when behavioral performance stabilizes.

In neural reassociation, the brain forms new stimulus-action associations by *reassociating* stimulus-code pairs instead of generating new activity patterns. Researchers observed neural reassociation in primate motor cortex during task learning [37]. NSM can reproduce neural reassociation with the correct motor function. Consider a nonlinear motor function *M* (**x**) from some unspecified brain region to motor cortex (instead of directly to an action). Let the function be nonlinear so that changing the distribution of inputs does not change the range of *M* (**x**), which we can think of an as a manifold in activity space. Suppose NSM operates only at the neurons in the domain of *M*. Then, task learning will occur by changing the structure of the neural activity in its domain to re-map stimuli to different but already existing patterns in motor cortex. If we were to record the neural activity in motor cortex during task learning, we would not observe significant changes in the manifold of response patterns. Instead, we would see the associations between these stimuli and the existing patterns shift. NSM can therefore reproduce neural reassociation.

We also discuss how the prediction of reward-dependent noise in the nervous system aligns with existing experimental evidence in Appendix I.

## VI. CONCLUSION AND OUTLOOK

This work proposes Neural Stochastic Modulation (NSM), a minimal and biologically plausible framework for unifying representation and policy learning in the brain. We focus on the core principles of the theory, leaving extensions and more complex tasks for future work. These results suggest further exploration into reward-dependent noise in the nervous system.

We outline several limitations and directions for future research, particularly regarding the efficiency of policy learning, in Appendix A.

## Appendix A Limitations & future directions

The present formulation of NSM is primarily limited by slow mixing in high-dimensional spaces, where its stochastic dynamics sample the steady-state distribution over weight space inefficiently. For example, imagine scaling the network’s representation space to *N* dimensions and placing it in a contextual bandit task with *N* actions and *N* input clusters. The time required for the network to discover an optimal policy by sampling from its limiting distribution scales poorly with *N*, as sampling occurs via a diffusive search across a high-dimensional sphere.

There are two potential solutions to this issue. The first is a biological argument that downplays it, and the second presents a few avenues for addressing the problem. Biological neural codes tend to be low dimensional [21, 38, 39]. If the weight dynamics are restricted to confine the activities largely to this low-dimensional subspace, then we avoid the poor scaling that comes with an increasingly high-dimensional weight space.

While plausible, this argument sidesteps the underlying issue. A more direct fix would be to make NSM’s sampling dynamics more efficient. The path forward becomes clearer when we distinguish between the two versions of NSM: one relying on indirect noise modulation and one on direct noise modulation.

In the indirect case, the network’s steady-state distribution explicitly balances reward and representation learning. The main challenge is efficient sampling, which is a classic problem in Markov Chain Monte Carlo methods. The NSM weight dynamics described here can be viewed as overdamped Langevin dynamics with position-dependent noise. Such dynamics can sample slowly when the target distribution is high-dimensional or non-log-concave, as may occur in our case. However, sampling efficiency can be improved. For example, underdamped Langevin dynamics [40] introduce momentum to better explore the distribution, while Riemannian Langevin dynamics [41], which would require nonuniform timescale modulation across the network, adapt to the geometry of parameter space. Both approaches could accelerate policy learning by improving convergence to the steady state.

In the case of direct noise modulation, NSM could exploit the full weight noise matrix ***σ***_*θ*_. In our examples, we assumed that neuromodulation acts uniformly across all neurons, so ***σ***_*θ*_ was proportional to the identity matrix. However, neuromodulation in biological systems is not uniform, which could allow for more structured and efficient diffusion. To see this, consider breaking up the agent’s interaction with the environment into “episodes”, and think of each episode as drawing a “sample” of the reward function *R* at a particular point in the network’s weight space. From this perspective, direct noise modulation often acts like an evolutionary algorithm operating over a constrained region of weight space– the solution space of ℒ. If the modulatory system shaped the noise matrix according to an evolutionary algorithm like Covariance Matrix Adaptation, it could accelerate policy learning by navigating the weight space more effectively.

A further promising direction is to integrate NSM with more expressive representation learning objectives, for instance, ones that construct predictive world models. Animals rapidly transfer knowledge between tasks by leveraging the common knowledge of how the world evolves under action. We include a very simple example of this knowledge transfer between tasks in Appendix J. Future work could explore whether agents with expressive representation learning objectives that build world models can solve more challenging tasks and generalize between them. It would also be interesting to explore other applications of the NSM framework: for instance, online supervised learning.

## Appendix B Neural Stochastic Modulation as a more general framework

While the focus of this work is to introduce NSM for policy optimization, we feel that it is important to briefly introduce the framework in a more general setting. NSM is a framework for how the nervous system can satisfy two distinct objectives when the primary network’s dynamics are governed by only one. Fundamentally, NSM states that:

1. Nervous systems are noisy,
2. Some objective ℒ determines the network’s deterministic weight dynamics,
3. A neuromodulatory system shapes the noise given the value of another objective *R*, either directly through modulating the weight noise matrix or indirectly by modulating the timescale.

The key insight behind NSM is that modulating the noise shapes the limiting distribution of learning, biasing the network towards configurations that optimize both ℒ and *R*. The objective ℒ does not need to be a representation learning objective, and *R* need not signify reward. The only requirement is that ℒ is differentiable with respect to the network weights. Nicely, we do not need to be able to compute the gradient of *R*.

Future work to explore other choices of ℒ or *R*. For instance, if *R* were a supervised learning loss instead of reward, the agent could learn a classification task online. There are a wealth of different objective combinations that one could consider.

We also note that one could further generalize the NSM framework outside of the context of neuroscience, to any model defined by parameters *θ* that aims to satisfy two objectives ℒ and *R*, though only ℒ is differentiable with respect to the model parameters. This framework may be useful if one believes that satisfying ℒ will help the model approach optimal solutions with respect to *R*.

## Appendix C Derivation of the stationary distribution for direct noise modulation

The goal of this section is to derive the stationary distribution of the network weights when learning is driven by direct noise modulation. We focus on the regime in which the noise magnitude is small enough that the weights remain confined to the solution space of the representation learning objective. This regime is particularly informative because it is analytically tractable and makes explicit how direct noise modulation can make reward a regularizer on representation learning.

We begin by recalling the general weight dynamics under direct noise modulation. Under this scheme, the net-work weights ***θ*** ∈ ℝ^*M*^ evolve according to the stochastic differential equation (SDE):

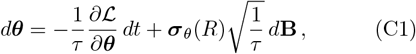

where:

- ***θ*** denotes the flattened weights of the network (a vector in ℝ^*M*^),
- *L*(***θ***) is a representation learning objective,
- *R*(***θ***) is the reward function,
- *τ >* 0 is a timescale constant,
- ***σ***_*θ*_(*R*) ∈ ℝ^*M×M*^ specifies reward-dependent noise statistics,
- *d***B** ∈ ℝ^*M*^ denotes a standard Wiener process (Brownian motion) in *M* dimensions.

We interpret this SDE in the Itô sense. The evolution of the probability density *p*(***θ***, *t*) is given by the corresponding Fokker–Planck equation:

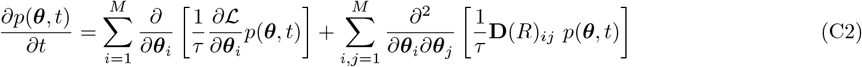

where 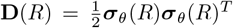 is the diffusion matrix. Subscripts denote elements on the corresponding vector or matrix.

In general, this equation does not yield an analytically tractable stationary distribution. However, since our goal is only to show that direct noise modulation makes reward act as a regularizer on representation learning, we consider a special case. Suppose that the noise matrix is isotropic,

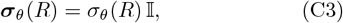

with

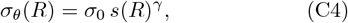

where *s*(*R*) is a monotonically decreasing function of *R, σ*_0_ *>* 0 is a baseline noise level, and *γ >* 0 a scaling constant. Noise decreases with greater reward.

Under this assumption, the Fokker–Planck equation becomes:

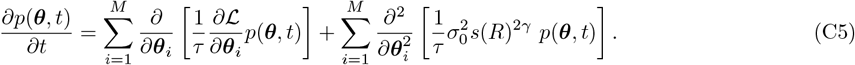

Next, we assume that the noise magnitude is sufficiently small for the weights to rapidly converge to, and remain near, the solution space of the representation learning objective ℒ. In this region ℛ ⊂ ℝ^*M*^, we have

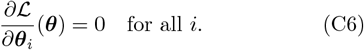

Within ℛ, the Fokker–Planck equation reduces to:

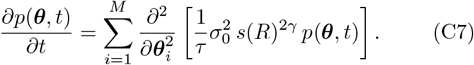

Since the weights are confined to ℛ, we must also impose reflecting (specifically, Neumann) boundary conditions on the boundary of this region. Setting the time derivative to zero yields the stationary distribution:

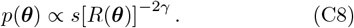

Equivalently, this can be expressed as:

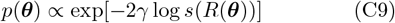

when ***θ*** ∈ ℛ and zero otherwise. One can verify that *p*(***θ***) satisfies the Neumann boundary condition. This distribution is identical to that of a system performing gradient descent on the effective objective

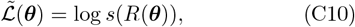

under fixed synaptic noise *γ*^−1*/*2^ within the solution space ℛ. Direct noise modulation makes reward act like a regularizer on representation learning, choosing solutions that maximize reward under the constraint that they are also optimal with respect to ∈ ℛ.

## Appendix D Derivation of the stationary distribution for indirect noise modulation

In this section, we derive the stationary distribution of network weights when learning occurs under indirect noise modulation.

Recall that under this scheme, the dynamics of the network weights ***θ*** ∈ ℝ^*M*^ are governed by the stochastic differential equation:

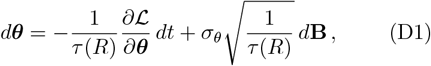

where the notation follows Appendix C. To understand the distribution of ***θ*** over time, we interpret the SDE in the Itô sense and write down the Fokker-Planck equation that describes the evolution of the probability density *p*(***θ***, *t*) over time:

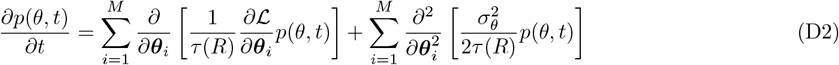

Note that we used the fact that the noise matrix is a multiple of the identity.

Setting the left hand side of the equation to zero gives the condition for the stationary distribution *p*(***θ***):

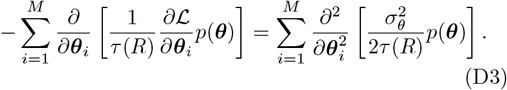

To proceed, we introduce a new function 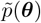:

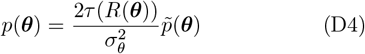

and plug this into Eq. (D3). This yields a simplified expression for 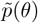

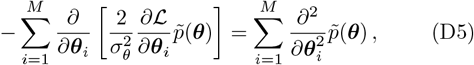

which has a Boltzmann form solution:

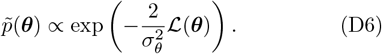

So, we have found that the stationary distribution of the learning dynamics is given by

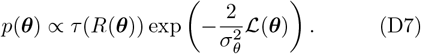

If we let

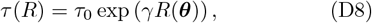

as required in Eq. (14), then we find

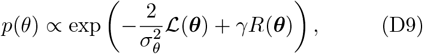

as claimed in the main text.

## Appendix E The Jacobian factor 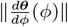 is independent of *ϕ*

In section IV.A.3 of the main text, we claimed that the Jacobian factor 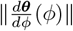 is independent of *ϕ*. We confirm this claim here.

In our example network, we have that ***θ*** = (**W**_0_, **W**_1_), where **W**_0_ ∈ ℝ^2*×D*^ represents feedforward connections from *D*-dimensional input states **s** ∈ ℝ^*D*^, while **W**_1_ ∈ ℝ^2*×*2^ represents recurrent lateral (inhibitory) connections within the network.

Recall from section IV.A.2 that the weights in the solution space of the representation learning object can be parameterized by a single angle *ϕ*. If we denote the rotation matrices in two dimensions as **U**(*ϕ*), then the set of weights in the solution space of the representation learning objective are

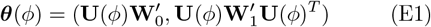

where 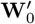 and 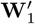 are chosen such that ***θ***(*ϕ* = 0) is in the solution space.

We can compute 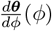 as:

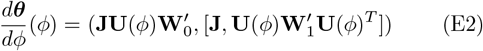

where

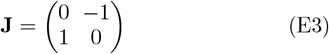

and 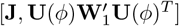 denotes the commutator of the two matrices. We can then compute

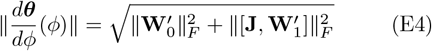

where ∥. ∥ is the Frobenius norm. So the Jacobian factor is independent of *ϕ*, as claimed in the main text.

## Appendix F Empirical support for NSM’s generality

The empirical investigations in the main text focus on a single example agent. We made this choice for clarity of presentation. However, it restricted the simulations that we discussed to one network architecture, one motor function, and one representation learning objective.

Our analytical results suggest generality across architectures and representation learning objectives. In deriving them, we made no reference to a particular choice of network architecture, motor function, or representation learning objective. This section decreases the gap between the generality of our analytical results and the lack of generality in our empirical investigations. We show that an NSM agent with a different network architecture and representation learning objective can learn rewarding policies in Acrobot, a more difficult control task than Cart Pole.

### 1. The agent has a neural network with an autoencoder architecture and a reconstruction loss

We consider a neural network with a linear autoencoder architecture. The network has three layers. The input and output layers are six-dimensional (this choice matches the observation space of Acrobot), while the hidden layer has three neurons. We let the hidden layer have three neurons because Acrobot has a discrete action space with three possibilities. This means that we can define the motor function on the hidden layer as an argmax over the hidden layer activities (i.e., each hidden neuron corresponds to a different action which the agent selects when this neuron is firing maximally). The representation learning objective is a standard reconstruction loss between the input and output of the network.

### 2. The training process follows indirect noise modulation

The learning process follows the indirect noise modulation protocol described in the main text. At each timestep of the task, the network activities converge to a feedforward pass on the observation. The network takes an action depending on which neuron in its hidden layer is maximally firing. The network then receives another observation, and the process continues. Throughout, the network’s weights perform noisy gradient descent on the reconstruction loss. A modulatory system alters the timescale of this noisy gradient descent based on a reward signal, according to Eq. (14) in the main text. Policy optimization emerges as a result of this modulation. We detail the specific parameters that we use during learning in Appendix M. We also made a few alterations to the process to stabilize learning, which we describe and justify below.

#### Timescale separation

Because the statistical structure of Acrobot observations is more complicated than those considered in the main text, we introduce a separation in timescales between the weight dynamics and the environmental dynamics. This choice stabilizes representation learning. In particular, we let the agent update its weights after every *P* episodes of interaction with the Acrobot task. We define an episode as a continuous sequence of timesteps, after which the double pendulum is reset due to success or failure. This timescale separation allows the network to estimate the reconstruction loss over order 500*P* timesteps (because there are at most 500 timesteps in an episode), which makes representation learning more robust. In our examples, we chose *P* = 150. This value is likely significantly larger than needed; we did not optimize it at all for efficiency. We refer to this engineering choice as a “separation of timescales” because, if we were to implement the same procedure in continuous time, it would amount to introducing a separation of timescales between weight and environmental dynamics. This separation of timescales in biologically realistic: Researchers have long observed synaptic plasticity in real brains evolve on the scale of minutes, which can be much longer than the timescale on which we receive novel stimuli from the environment [42].

#### Reward structure adjustment

We also tweak the Acrobot reward structure to facilitate policy learning. The traditional Acrobot reward structure gives the agent a reward of −1 during each timestep that it has not yet swung the double pendulum above the target height. The episode stops either when the agent succeeds in reaching the target or after 500 timesteps. So, after every episode, the agent has received a reward *R*_traditional_ equal to negative the number of timesteps it takes to reach the target height, capped at −500. Since the weights change on the timescale of episodes rather than individual timesteps, only this episode-based reward structure affects policy learning performance. We use the subscript “traditional” to denote this typical reward structure. We change the Acrobot episode-based reward structure in the following manner. We give the agent *R*_adjusted_ = 0 if the traditional Acrobot reward structure specified *R*_traditional_ = −500 at the end of an episode. This is a simple baseline shift that is equivalent to changing our agent’s hyperparameters. However, we also give the agent *R*_adjusted_ = 200 if it receives traditional reward *R*_traditional_ *>* −500+*J*, where *J* is a threshold hyperparameter over which we sweep. This change enforces that the agent only receives reward when it performs the task well enough, where “well enough” is specified by the threshold *J*.

The altered reward structure biases the effective loss only towards weight configurations the lead to policies with *R*_traditional_ *>* −500 + *J*, encouraging the agent to not only get the double pendulum to the target, but also to succeed sufficiently rapidly. In principle, one could get the same results by increasing the strength of timescale modulation *γ* (see Eq. 16 in the main text). However, because we have more intuition for the effect of specific values of *J* than the effect specific values of *γ* on the performance of the agent, with limited compute resources, we found that hyperparameter searching was easier when we searched over *J* because we knew how to initialize at reasonable values and define reasonable ranges a priori.

### 3. The agent learns to beat Acrobot

After a hyperparameter sweep over a subset of the parameters (*σ*_*θ*_, *J*), we found a region in hyperparameter space for which the NSM agents consistently beat or nearly beat Acrobot (Fig 5). We note that the agent’s performance is fairly robust to the specific choice of hyperparameters.

**FIG. 5.**
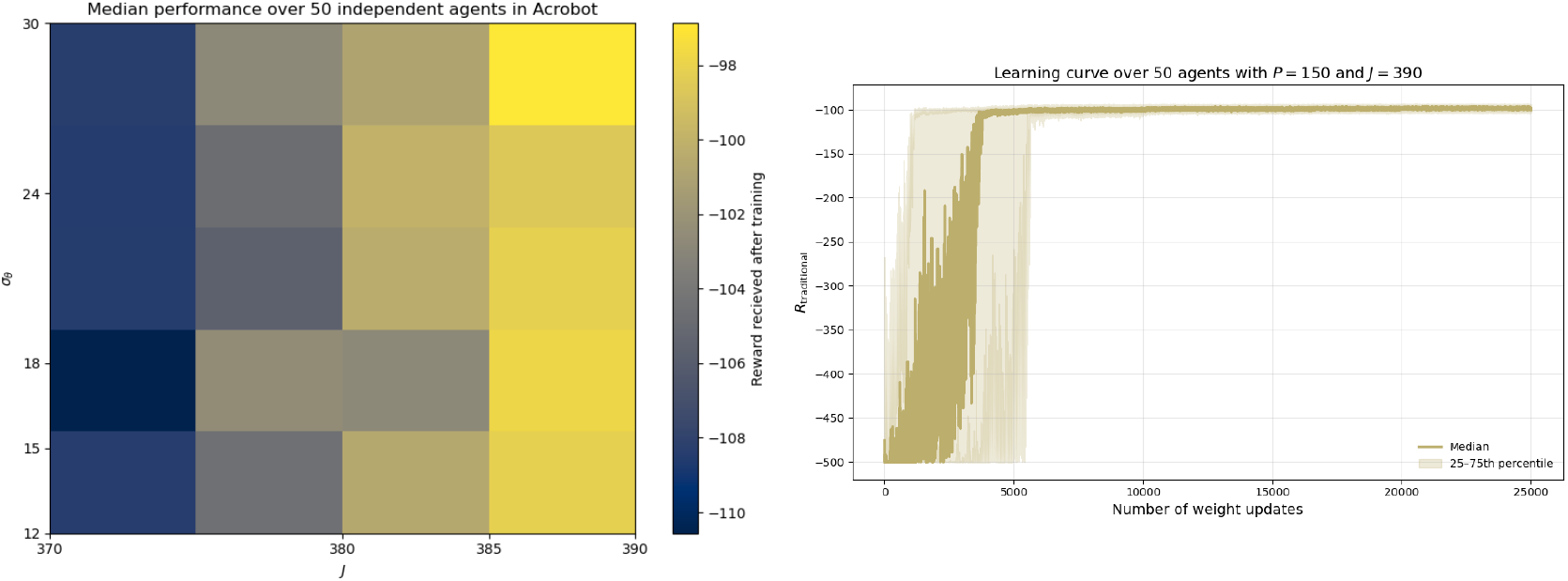
**(Left)** Median performance across 50 independent agents for different hyperparameter settings. Color indicates episode reward in standard Acrobot units (*R*_traditional_). The x-axis varies *J*, the threshold on *R*_traditional_ above which the agent receives a reward signal, while the y-axis varies *σ*_*θ*_, the baseline standard deviation of weight noise. A median reward above −100 is considered beating Acrobot, and values above −500 correspond to successfully reaching the target height. The agent achieves near- or above-baseline performance across a range of hyperparameter. **(Right)** Learning curves for 50 agents using the hyperparameters from the upper-right corner of the left panel. The dark line shows the median reward; the shaded region denotes the 25th–75th percentile range. Agents beat Acrobot after roughly 3,000 weight updates and maintain stable performance thereafter.

We also present the learning curve for the best set of hyperparameters in Fig. 5. We find that agents typically learn the task after about 3, 000 weight updates. We choose to measure learning time in weight updates rather than environmental timesteps because we are modeling a real agent that lives in continuous time. For such an agent, the relevant timescale over which to measure learning speed is the timescale over which the weights change. In discretized time, as we have here, the learning time in units of the weight timescale is the proportional to the number of weight updates.

NSM’s success with a different network architecture, representation learning objective, and motor function provides numerical support for NSM’s generality. Given that Acrobot is also a more difficult control task, these experiments also show NSM’s promise for scaling to more difficult settings, despite the concerns highlighted in Appendix A regarding slow mixing in high dimensions.

## Appendix G The network weights remain in the solution space of ℒ during the contextual bandit task

Here we provide empirical evidence of our claim that the weights of the representation learning network converge to and remain within the solution space of ℒ during the context bandit task. This claim is important because it allows us to describe the network weights with a single order parameter *ϕ* and apply our general results on direct noise modulation.

In a general task with reward-dependent noise, we cannot guarantee that weights converge to the solution space of ℒ. As discussed in the main text, reward-dependent noise can stabilize points outside of the solution space if it improves the agent’s policy.

However, we designed the contextual bandit task so that the two Gaussian clusters are only linearly separable along the second principal component of the inputs. This choice ensures that optimal policies exist within the solution space of ℒ, which prevents the weights from settling outside it.

Figure 6 empirically confirms that the principal subspace error rapidly decreases and remains near optimal throughout this task. The weights remain approximately within the solution space of ℒ, as claimed in the main text.

**FIG. 6.**
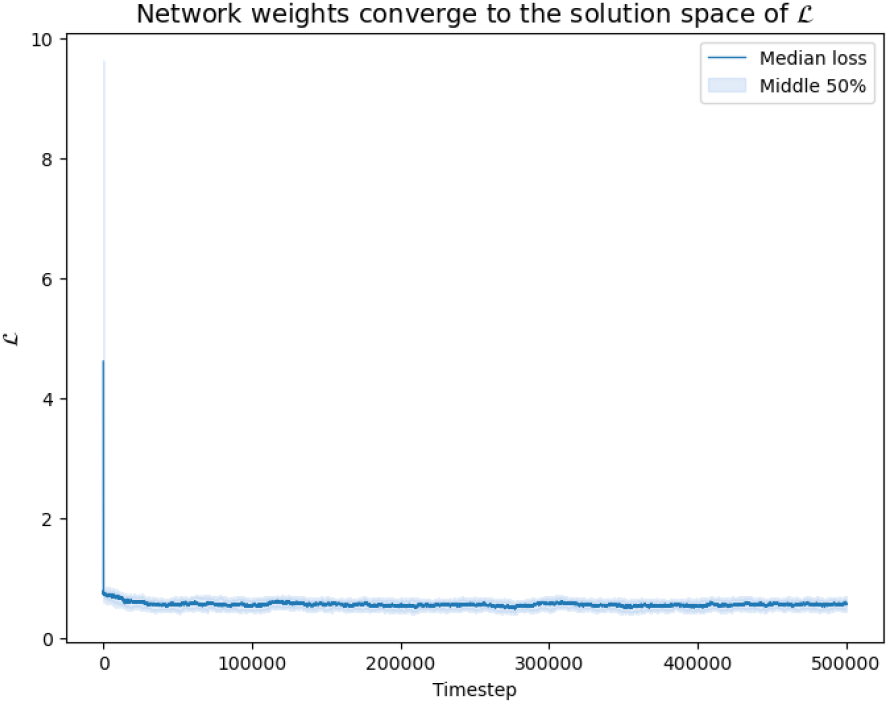
The representation learning loss, quantified by 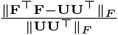. The columns of **U** are the top two eigenvectors of the covariance matrix **C**, and 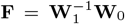 is the projection performed by the network after the convergence of (4). The operation ∥ · ∥_*F*_ denotes the Frobenius norm.

## Appendix H The role of activity noise in NSM policy optimization

This work shows that reward-dependent *weight* noise facilitates policy optimization in networks otherwise designed to learn representations. It is natural to ask: Can activity noise play the same role? Or is its only effect to make the agent’s policy stochastic? Here, we answer this question by showing that activity noise is an “effective” weight noise, and therefore, the network can harness it for policy optimization. Intuitively, activity noise acts as an effective weight noise because the weight updates depend on the activities. We show this in the case of the Similarity Matching objective for concreteness. We consider discrete timesteps throughout. However, it will be clear that the argument applies to any representation learning objective and to continuous time.

Consider the two-neuron network presented in this work. Recall that it has activities **x** ∈ ℝ^2^ and weights *θ* = (**W**_0_, **W**_1_). Let **x**_0_(**s**) denote the network’s response to a stimulus **s** without the presence of noise. Then the network’s response with noise **x** is given by

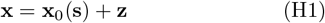

where **z** ∈ ℝ^2^ is a random variable with covariance matrix 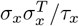, following Eq. (1). Per Eqs. (5-6), considering discrete timesteps and setting the weight noise to zero, the weight updates are given by

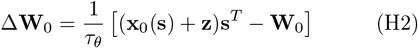

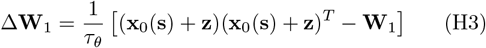

which can be expanded into

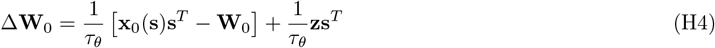

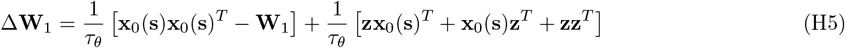

We recognize that the first term in each equation corresponds to the weight updates without activity noise. The second terms are effective weight noise terms; they depend on **z**. As such, NSM policy learning can also occur through activity noise (or some combination of activity and weight noise). Note that the learning rate 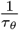 modulates this effective weight noise, as it does the pure weight noise (through the timescale of the stochastic dynamics). This is especially interesting given evidence that dopamine, a neuromodulator associated with reward signals, has been observed to tune the weight learning rate (i.e., timescale) and the activity signal-to-noise ratio in certain regions of mammalian nervous systems [43–46].

We further emphasize that a system can in principle implement both direct noise modulation through the activity noise statistics and indirect noise modulation through the timescale of the weight dynamics using only this “effective” weight noise.

## APPENDIX I The biological plausibility of the NSM noise hypothesis

Here, we discuss the biological plausibility of the NSM reward-dependent noise hypothesis. Is there any evidence that the brain uses reward-dependent noise? Directly measuring weight noise is experimentally challenging because it is difficult to precisely track synaptic weight dynamics. However, we can look for indirect evidence.

Recall that, in indirect noise modulation, NSM tunes the network’s weight noise through the timescale of the stochastic weight dynamics. This aligns with findings showing that dopamine, a neuromodulator associated with reward signals, adapts the timescale of the weight dynamics in mice during policy optimization [43]. (Note, however, that the authors refer to this “timescale” as a “learning rate,” which is a term that we use to refer to a related but different quantity.)

Additionally, in Appendix H, we find that activity noise can serve as an effective weight noise to guide policy learning. Research suggests that dopamine also tunes the activity signal-to-noise ratio in certain regions of mammalian nervous systems [44–46], equivalent to modulating activity noise. Together, these examples suggest that dopamine may adapt synaptic noise in response to reward signals, supporting the NSM reward hypothesis. However, future research is needed to investigate this claim in greater detail.

The dopaminergic system is not the only component of the nervous system connected to both synaptic noise and reward signals. There is also evidence linking astrocytes, a type of glial cell that makes up the majority of cells in the central nervous system, to synaptic noise modulation [47] and reward-seeking behavior [48]. Again, more research is needed to investigate whether these connections are coincidental or a signature of NSM at work.

## Appendix J NSM exhibits efficient knowledge transfer across tasks

Organisms do not relearn the dynamics of their environment each time they face a new task. Instead, they transfer their knowledge from past experiences, using previously learned representations to learn new sequences of actions. NSM agents demonstrate similar knowledge transfer across tasks in online learning settings. Specifically, when a task switch occurs, NSM agents leverage the information already encoded in their representation learning networks to adapt and optimize a new policy.

We illustrate this idea with a modified version of the contextual bandit task considered in Section 4.1. Previously, the agent needed to learn to take the action *a* corresponding to the random sign *k*. In the modified task, the rewarding action-sign association changes from *a* = *k* to *a* = −*k* on a timescale unknown to the agent. Each association represents a different task. Figure 7 (right) illustrates that the agent adapts to the changing tasks. Importantly, the agent does need to relearn to extract the principal subspace of the inputs (Figure 7, left). Instead, it only needs to relearn how to use this information. Knowledge is transferred efficiently as the tasks evolve.

**FIG. 7.**
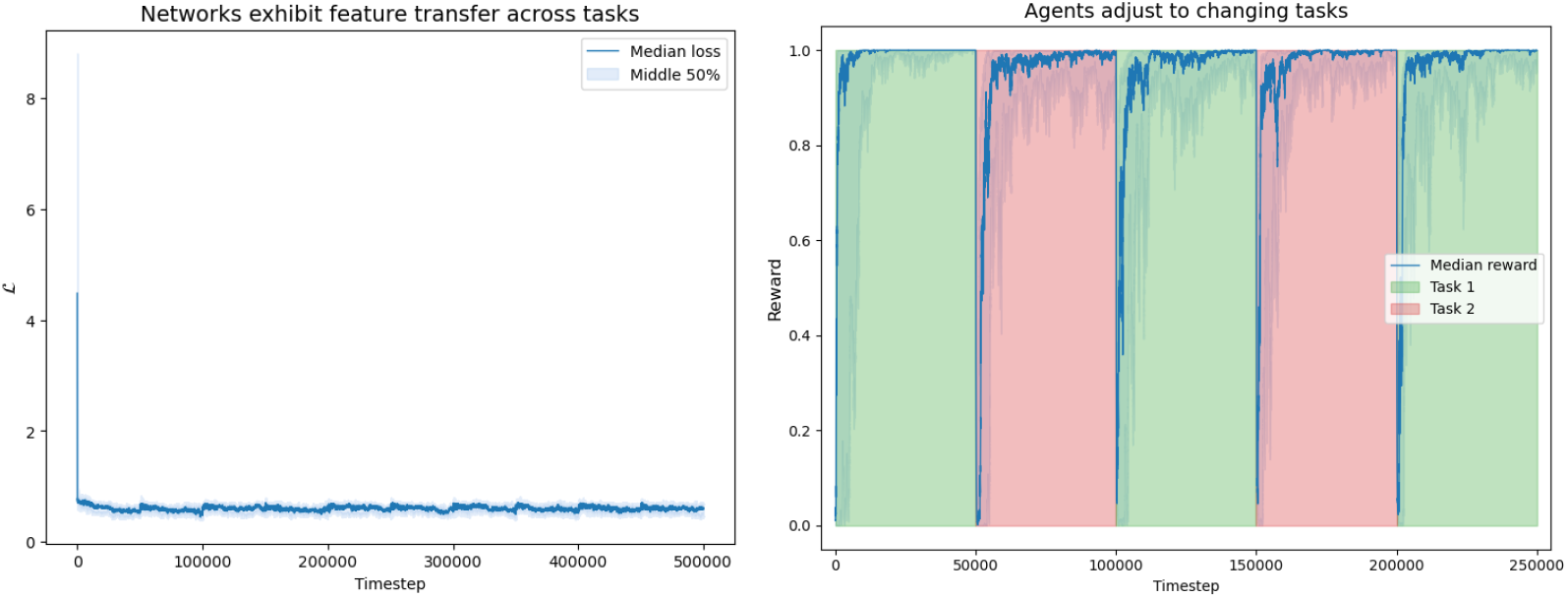
(Left) The principal subspace loss remains low as tasks change, facilitating efficient knowledge transfer. (Right) The agent adjusts readily to changing tasks.

The NSM agents we focus on in this work quickly optimize their representation learning objective. As a result, transferring learned features between tasks doesn’t significantly affect their efficiency. However, for agents with more complex representation learning objectives, this transfer becomes crucial. It allows agents to learn the dynamics of the environment just once, after which they can use this understanding to solve a variety of tasks.

## Appendix K Approximation of *R*(*ϕ*)

In this section, we approximate *R*(*ϕ*) as a function of the contextual bandit state distribution. This calculation is necessary to write down the stationary distribution over *ϕ* during learning, which tells us about which policies the agent tends to learn. Our strategy will be as follows:

1. We will write *a*_*ϕ*_(**s**), the agent’s action under an observation **s** as a function of the angle *ϕ*.
2. We will write *R*(*ϕ*) as an expectation value and eventually an integral.
3. We will see that evaluating this expression requires finding the distribution of the network responses **x**(**s**). To find this distribution, we compute the principal subspace of **s**.
4. We then use this information to write the distribution of **x** as a function of *ϕ*.
5. Finally, we approximate the integral in step (2).

### 2. Step 1: Finding *a*_*ϕ*_(**s**)

We begin by finding *a*_*ϕ*_(**s**). As in the main text, we assume that the agent’s parameters are in the solution space of ℒ. It follows that its activity dynamics (4) that its parameters *θ* define a mapping from the observation space to the network’s representation space given by

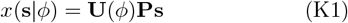

where the operator *P* projects **s** into its 2-dimensional principal subspace, and **U**(*ϕ*) ∈ ℝ^2*×*2^ rotates the result by *ϕ* radians. By convention, we choose **P** such that **x**(*k****µ*** | *ϕ* = 0) = *ck***v** for some *c >* 0.

Since *M* (**x**) = sign(**v**^*T*^ **x**), with 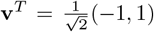, this corresponds to the action

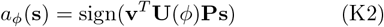

### 2. Step 2: Write *R*(*ϕ*) **as an integral**

Recall that the goal of the agent is to pick the action *k*, the random sign. Then the reward at a single step, *r*(**s**, *a*), is given by

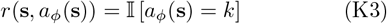

Since the distribution of **s** is stationary and the agent’s policy is fixed, the expected reward 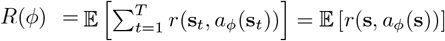. So, we can write *R*(*ϕ*) as

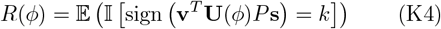

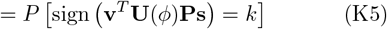

Let *f*_**x**_(**x**_1_, **x**_2_ | *k, ϕ*) be the distribution of the network representations **x** conditional on the random sign *k* and given *ϕ*. Then, considering the meaning of (24) geometrically, we see that is it equivalent to

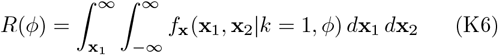

We aim to evaluate this integral.

### 3. Step 3: Finding the principal subspace of s

To evaluate the integral (25), we must find the distribution of **x** = *U* (*ϕ*)*P* **s**. Of course, to find this distribution we must first determine *P*. So, we need to find the principal subspace of **s**. Recall that we can decompose 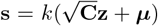, where **z** is a standard normal random variable in 10 dimensions, **C** is the covariance matrix, ***µ*** is the mean, and k is a random sign. Since **s** is centered (due to the random sign), its covariance matrix is given by

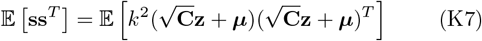

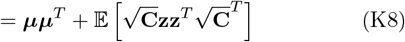

where we used the fact that · [**z**] = 0. Then, using · **zz**^*T*^ = 𝕀, and since **C** is symmetric, 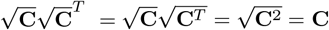, we find

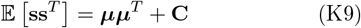

In the contextual bandit task, we defined **C** such that ***µ*** is an eigenvector. Let *λ*_***µ***_ be the eigenvalue associated with ***µ***, and 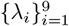 denote the other nine eigenvalues with associated eigenvectors 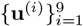. Then the eigenvalues of · **ss**^*T*^ are given by {∥***µ***∥^2^ + *λ*_***µ***_, *λ*_1_, …, *λ*_9_}.

The mean vector ***µ*** allows the agent to identify the random sign *k*. So, to ensure that there exists a weight configuration in the solution space of ℒ that solves the task, we needed the projection of ***µ*** onto the principal subspace of **s** to be nonzero. We therefore chose ∥***µ***∥^2^ + *λ*_***µ***_ *> λ*_1_ *>* … ≥ *λ*_9_.

We see that the principal subspace of **s** is given by span(***µ*, u**^(1)^). We can therefore define **P** as the projection onto this subspace with *P* ***µ*** = ∥***µ***∥**v** and 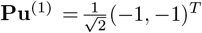.

### 4. Step 4: Finding the distribution *f*_**x**_(**x**_1_, **x**_2_|*k* = 1, *ϕ*)

Now that we have determined the projection operator **P**, we can find the distribution of **x** given the random sign *k* = 1. Notice that, conditioned on *k*, the distribution of **s** is Multivariate Normal. Any projection or rotation of a Multivariate Normal is still Multivariate Normal. Nicely, this means that the distribution of **x**(*s*) = **U**(*ϕ*)**Ps** is defined by its mean and covariance matrix. We need only compute these.

Recall that 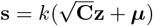, so

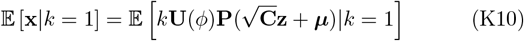

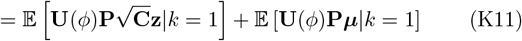

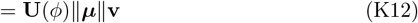

where, in the last line, we used *P* ***µ*** = ∥***µ*** ∥ **v** and · [**z**] = 0. Before we evaluate the covariance matrix, we make the following simplification. Let ***µ***′ := <u>∥</u>***<u>µ</u>***∥**v**. Notice that

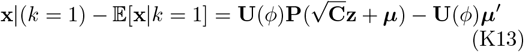

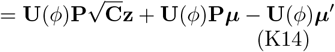

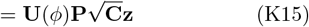

We can now evaluate the covariance matrix, which we denote **C**′.

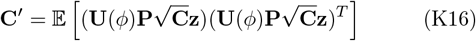

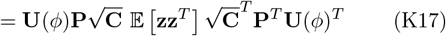

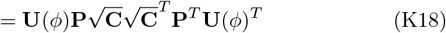

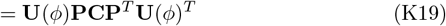

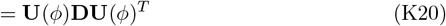

where **D** = diag(*λ*_***µ***_, *λ*_1_). We made the last step by the definition of the projection operator **P**.

This means that

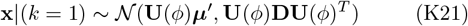

We can now attempt to evaluate the integral (47).

#### a. Step 5: Approximating the integral for R(ϕ)

We now see that

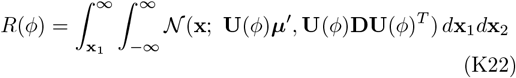

This integral is not analytically tractable because the Gaussian is not centered and has a potentially nondiagonal covariance matrix. However, we can make an approximation that will be valid for our contextual bandit task.

Notice that we defined the contextual bandit state distribution such that *λ*_***µ***_ *<<* ∥***µ***∥ ^2^, *λ*_1_. We did this so that the agent can learn to solve the task consistently (the two Gaussian clusters need to be linearly distinguishable). We therefore approximate the expression with the limit that *λ*_***µ***_ → 0.

For geometric intuition, refer to Figure 8. We see that, in this limit, the integral would be exactly given by

**FIG. 8.**
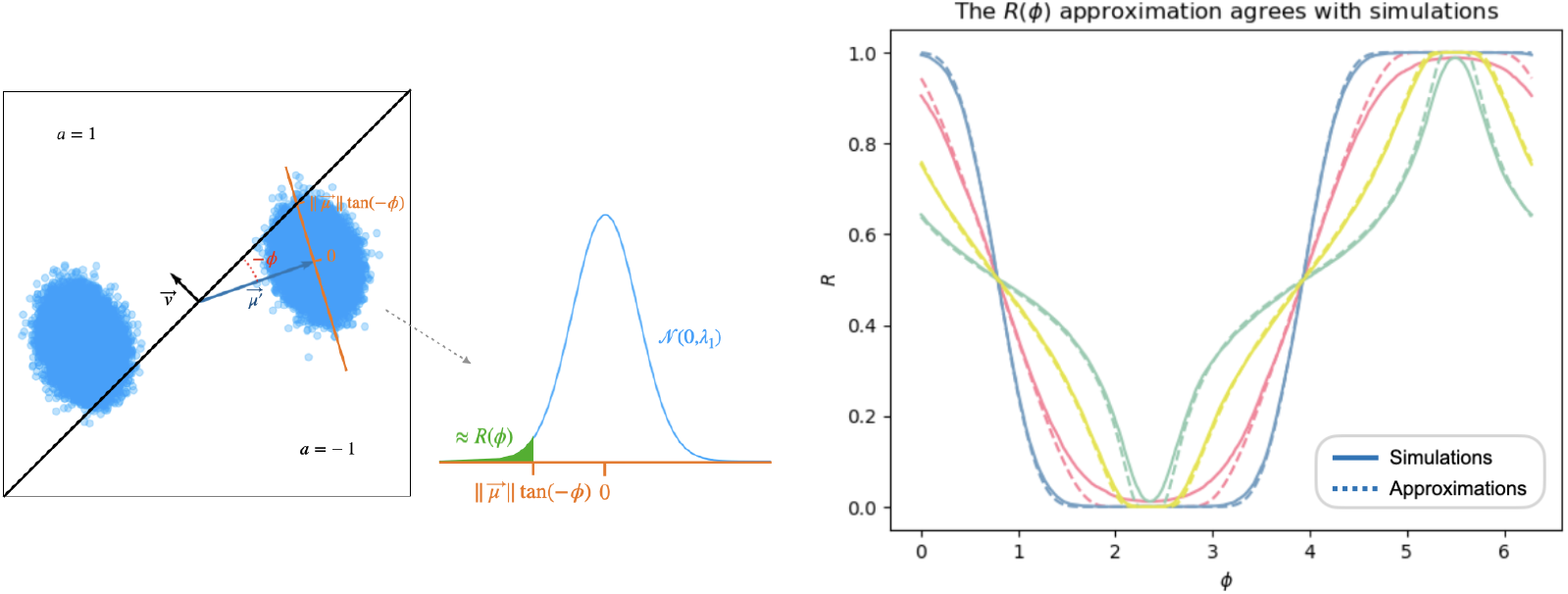
**(Left)** A visualization of the *R*(*ϕ*) approximation. *R*(*ϕ*) is equivalent to the probability that the activity pattern **x**(**s**|*ϕ, k* = 1) is mapped by the motor function to *a* = 1. Visually, this corresponds to the probability that the rightmost cluster is in the upper region of activity space. In the limit that *λ*_***µ***_ → 0, all variance perpendicular to the orange line disappears. So this quantity becomes the integral of the distribution 𝒩 (0, *λ*_1_) from −∞ to |***µ***∥ tan(*ϕ*). **(Right)** The *R*(*ϕ*) approximation is reasonable across a range of examples where our assumption *λ*_***µ***_ *<<* ∥***µ***∥^2^, *λ*_1_ holds. Solid lines indicate simulation values, and dashed lines denote the analytical approximation. In these simulations, ∥***µ***∥^2^ ∈ {0.25, 1}, *λ*_1_ ∈ {0.1, 2}, and *λ*_***µ***_ = 0.05.

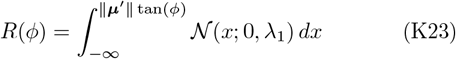

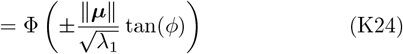

since ∥***µ***′∥ = ∥***µ***∥, where we take the negative sign when 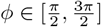 and the positive sign otherwise.

### 5. Our approximation agrees with simulations

In Figure 8, we verify our approximation by comparing it to simulations. We find good agreement when our assumption that *λ*_*µ*_ *<<*∥ ***µ*** ∥^2^, *λ*_1_ holds. To get simulation values of *R*(*ϕ*), we define the mapping *a*_*ϕ*_(**s**). We then draw the states **s** according to the process defined in the main text and compute the expected value of the indicator that *a*_*ϕ*_(**s**) = *k*, where *k* is the random sign.

## Appendix L Optimal Cart Pole policies do not exist in the solution space of ℒ

Here, we show that optimal Cart Pole policies cannot be found within the solution space of the Similarity Matching objective ℒ. We begin with an intuitive argument. Suppose, for the sake of contradiction, that the agent can maintain a successful policy while satisfying ℒ. A successful Cart Pole policy minimizes the angular deviation of the pole from its center. However, because the agent satisfies ℒ, it projects its observations onto their two-dimensional principal subspace. The pole’s angular deviation does not lie in this subspace, as its variance is too small. Without access to this critical information, the agent cannot determine which action to take, and thus cannot have an optimal policy. This leads to a contradiction.

It is necessary to validate this intuition because the other observations (cart position, velocity, etc.) should have small variances in an optimal policy as well. We test the claim that an agent with an optimal policy cannot project its observations onto their 2D principal subspace and maintain an optimal policy. We record the observations of an agent with a policy that beats Cart Pole and compute their principal subspace. We then define a new agent with a motor function *M* (**x**) = sign(**v**^*T*^ **x**) that acts directly on these principal subspace projections. We explore all rotational configurations of the principle subspace projections but find that none result in a policy that beats Cart Pole (Figure 9). An agent cannot project its observations onto their principal subspace if it intends to maintain its optimal policy^7^. We conclude that agents cannot simultaneously exist in the solution space of ℒ and have an optimal policy. This result empirically confirms the claim that optimal policies do not exist within the solution space of ℒ.

**FIG. 9.**
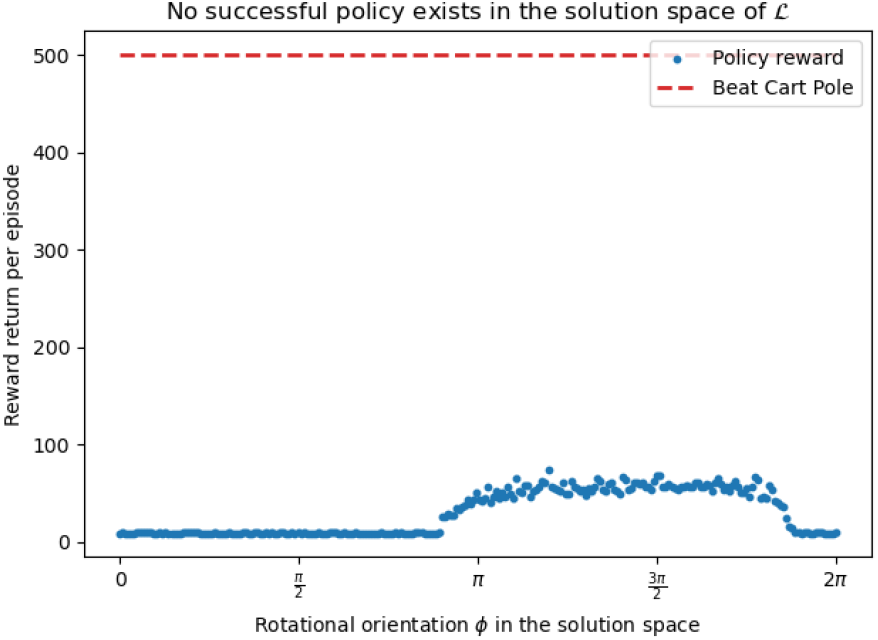
No rotational orientation of the network activities can solve Cart Pole if the observations are projected onto their principal subspace. Points give the median reward return for an agent with a given rotational orientation. *ϕ* is the same angle depicted in Figure 2.

## Appendix M Agent parameters in the three tasks

Table 1 provides the parameters that we used to run the agent simulations in the contextual bandit and Cart Pole tasks. We note here that we did not add noise to **W**_1_ during the simulations. We made this choice because the SM objective requires that **W**_1_ remain symmetric. Alternatively, we could have chosen to symmetrize this weight noise, which would have given similar results. The Acrobot task had noise on all weights.

**TABLE 1.** Agent Parameters.

| Task | $\gamma$ | $\sigma_\theta$ | $\tau_\theta$ |
| --- | --- | --- | --- |
| **Contextual Bandit** | $\{0, 1/2, 1, 3/2, 2, 5/2\}$ | $0.001 \exp(-\gamma R)$ | 830 |
| **Cart Pole** | 1/30 | 1 | $100 \exp(\gamma R)$ |
| **Acrobot** | 1 | $\{12, 15, 18, 24, 30\}$ | $1000 \exp(\gamma R)$ |

## Appendix N Implementation of the modulatory system *R* estimates

In this article, we often refer to a modulatory system that estimates *R*. However, we did not mention how this system implements this policy reward estimation.

### 1. The contextual bandit

In the contextual bandit task, the modulatory system estimates the reward *R* with an exponential filter of length *T* = 100 timesteps. It updates its reward esti-mate at time 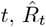, with the equation

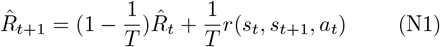

where *r*(*s*_*t*_, *s*_*t*+1_, *a*_*t*_) gives the reward received after that timestep.

### 2. Cart Pole

In Cart Pole, the modulatory system uses the last trial result as the *R* estimate. This is equivalent to the contextual bandit implementation if we consider the exponential filter to be over trials instead of timesteps, and we set *T* = 1.

### 3. Acrobot

In Acrobot, the modulatory system estimates reward differently. Rather than an exponential filter, it uses the maximum value received over the previous *T* Acrobot episodes. While this choice overestimates the true average reward produced by the agent’s policy, we empirically find that it stabilizes policy learning by producing lower variance in the reward estimates. It makes sense that this estimator has lower variance than an exponential filter because, typically, the largest order statistic from *N* samples of a distribution that is bounded from above (for example, a sample from the reward received in an Acrobot trial) has lower variance than the average of these samples. We a bias in the modulatory system’s reward estimates to lower their variance.

When reporting agent performance, we still report the sample mean over independent trials, not the maximum.

## Footnotes

1 We define a Markov Decision Process (MDP) by the tuple (𝒮, 𝒜′, P, r, γ), where 𝒮 is the state space, 𝒜 is the action space, P (s′ |′s, a) gives the probability of transitioning from the state s to s′ under the action a, the function r(s, s′, a) assigns a re-ward to this event, and γ ∈ (0, 1) is the discount factor. The agent follows a policy π, such that at time t its action is sampled from the distribution a_t_ ~ π(a | s_t_). The agent aims to learn a policy which maximizes the expected discounted reward 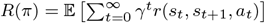.

2 **W**_1_ must be symmetric and positive-definite.

3 This is an assumption that is not necessary but will simplify the discussion.

4 A similar result has been found in the study of the coordinatedependent diffusion of particles [31].

5 The choice to make ***σ***_***θ***_ diagonal is an unnecessary simplification that may reduce policy learning efficiency for direct noise modulation (see Limitations).

6 In practice, convergence is much faster.

7 We assumed here that the result in Figure 9 hold for all successful policies, not just the ones tested. This assumption is reasonable because Cart Pole is a simple task; there are not successful policies that lead to drastically different input distributions.

## References

[1] R. P. Rao and D. H. Ballard, Predictive coding in the visual cortex: a functional interpretation of some extraclassical receptive-field effects, Nature neuroscience 2, 79 (1999).

[2] P. Perruchet and S. Pacton, Implicit learning and statistical learning: One phenomenon, two approaches, Trends in Cognitive Sciences 10, 233 (2006).

[3] E. L. Thorndike, Animal intelligence: An experimental study of the associative processes in animals, The Psychological Review: Monograph Supplements 2, i (1898).

[4] D. O. Hebb, The organization of behavior: A neuropsychological theory (Psychology Press, 2005).

[5] L. F. Abbott and S. B. Nelson, Synaptic plasticity: taming the beast, Nature neuroscience 3, 1178 (2000).

[6] T. Konkle and G. A. Alvarez, A self-supervised domain-general learning framework for human ventral stream representation, Nature Communications 13, 491 (2022).

[7] C. Zhuang et al., Unsupervised neural network models of the ventral visual stream, Proceedings of the National Academy of Sciences 118, e2014196118 (2021).

[8] M. S. Halvagal and F. Zenke, The combination of hebbian and predictive plasticity learns invariant object representations in deep sensory networks, Nature Neuroscience 26 (2023).

[9] A. Sengupta et al., Manifold-tiling localized receptive fields are optimal in similarity-preserving neural networks, in Advances in Neural Information Processing Systems, Vol. 31 (2018).

[10] S. Bakhtiari et al., The functional specialization of visual cortex emerges from training parallel pathways with self-supervised predictive learning, Advances in Neural Information Processing Systems (2021).

[11] W. Lotter, G. Kreiman, and D. Cox, A neural network trained for prediction mimics diverse features of biological neurons and perception, Nature Machine Intelligence 2, 210 (2020).

[12] R. V. Raju et al., Space is a latent sequence: A theory of the hippocampus, Science Advances 10, eadm8470 (2024).

[13] R. Schaeffer et al., Self-supervised learning of representations for space generates multi-modular grid cells, in Advances in Neural Information Processing Systems, Vol. 36 (2023) pp. 23140–23157.

[14] D. Levenstein et al., Sequential predictive learning is a unifying theory for hippocampal representation and replay (2024), bioRxiv: 2024–04.

[15] K. K. Nejad et al., Self-supervised predictive learning accounts for cortical layer-specificity, Nature Communications (2025).

[16] R. V. Florian, Reinforcement learning through modulation of spike-timing-dependent synaptic plasticity, Neural Computation 19, 1468 (2007).

[17] R. Legenstein, D. Pecevski, and W. Maass, A learning theory for reward-modulated spike-timing-dependent plasticity with application to biofeedback, PLoS Computational Biology 4, e1000180 (2008).

[18] H. Markram et al., Regulation of synaptic efficacy by coincidence of postsynaptic aps and epsps, Science 275, 213 (1997).

[19] G.-q. Bi and M.-m. Poo, Synaptic modifications in cultured hippocampal neurons: dependence on spike timing, synaptic strength, and postsynaptic cell type, Journal of Neuroscience 18, 10464 (1998).

[20] J. Qin and A. R. Wheeler, Maze exploration and learning in c. elegans, Lab on a Chip 7, 186 (2007).

[21] J. M. Shine et al., Human cognition involves the dynamic integration of neural activity and neuromodulatory systems, Nature Neuroscience 22, 289 (2019).

[22] M. Jaderberg et al., Reinforcement learning with unsupervised auxiliary tasks, arXiv preprint 1611.05397 (2016).

[23] D. Ha and J. Schmidhuber, Recurrent world models facilitate policy evolution, Advances in Neural Information Processing Systems 31 (2018).

[24] A. Greco, A. C. Schapiro, R. Koster, and C. Baldassano, Predictive learning shapes the representational geometry of the human brain, Nature Communications 15, 9670 (2024).

[25] A. Clark, Whatever next? predictive brains, situated agents, and the future of cognitive science, Behavioral and Brain Sciences 36, 181 (2013).

[26] C. Capone and P. S. Paolucci, Towards biologically plausible model-based reinforcement learning in recurrent spiking networks by dreaming new experiences, Scientific Reports 14, 14656 (2024).

[27] D. Kappel et al., Reward-based stochastic self-configuration of neural circuits, arXiv preprint 1704.04238, 1162 (2017).

[28] W. Maass, Noise as a resource for computation and learning in networks of spiking neurons, Proceedings of the IEEE 102, 860 (2014).

[29] C. Li et al., Neuron-level prediction and noise can implement flexible reward-seeking behavior, arXiv preprint (2024), bioRxiv: 2024-05.

[30] C. Pehlevan, T. Hu, and D. B. Chklovskii, A hebbian/anti-hebbian neural network for linear subspace learning: A derivation from multidimensional scaling of streaming data, Neural Computation 27, 1461 (2015).

[31] R. Maniar and A. R. I. J. I. T. Bhattacharyay, Random walk model for coordinate-dependent diffusion in a force field, Physica A: Statistical Mechanics and its Applications 584, 126348 (2021).

[32] S. Qin et al., Coordinated drift of receptive fields in hebbian/anti-hebbian network models during noisy representation learning, Nature Neuroscience 26, 339 (2023).

[33] F. Chen et al., Stochastic collapse: How gradient noise attracts sgd dynamics towards simpler subnetworks, Advances in Neural Information Processing Systems 36 (2024).

[34] L. N. Driscoll et al., Dynamic reorganization of neuronal activity patterns in parietal cortex, Cell 170, 986 (2017).

[35] A. Rubin et al., Hippocampal ensemble dynamics timestamp events in long-term memory, eLife 4, e12247 (2015).

[36] Y. Ziv et al., Long-term dynamics of ca1 hippocampal place codes, Nature Neuroscience 16, 264 (2013).

[37] M. D. Golub et al., Learning by neural reassociation, Nature Neuroscience 21, 607 (2018).

[38] K. K. Iyer et al., Focal neural perturbations reshape low-dimensional trajectories of brain activity supporting cognitive performance, Nature Communications 13, 4 (2022).

[39] R. B. Ebitz and B. Y. Hayden, The population doctrine revolution in cognitive neurophysiology, Neuron 109, 3055 (2021).

[40] X. Cheng et al., Underdamped langevin MCMC: A nonasymptotic analysis, in Conference on Learning Theory (PMLR, 2018) pp. 300–323.

[41] M. Girolami and B. Calderhead, Riemann manifold langevin and hamiltonian monte carlo methods, Journal of the Royal Statistical Society: Series B (Statistical Methodology) 73, 123 (2011).

[42] K. C. Bittner et al., Behavioral time scale synaptic plasticity underlies ca1 place fields, Science 357, 1033 (2017).

[43] L. T. Coddington, S. E. Lindo, and J. T. Dudman, Mesolimbic dopamine adapts the rate of learning from action, Nature 614, 294 (2023).

[44] C. M. Vander Weele et al., Dopamine enhances signalto-noise ratio in cortical-brainstem encoding of aversive stimuli, Nature 563, 397 (2018).

[45] G. Winterer and D. R. Weinberger, Genes, dopamine and cortical signal-to-noise ratio in schizophrenia, Trends in Neurosciences 27, 683 (2004).

[46] S. Kroener et al., Dopamine modulates persistent synaptic activity and enhances the signal-to-noise ratio in the prefrontal cortex, PloS One 4, e6507 (2009).

[47] S. Lewis, Astrocytes make synapses noisy, Nature Reviews Neuroscience 13, 451 (2012).

[48] S. Kang et al., Astrocyte activities in the external globus pallidus regulate action-selection strategies in reward-seeking behaviors, Science Advances 9, eadh9239 (2023).

